# Odors Smell Like Their Components: A Linear Framework for Predicting Olfactory Mixture Perception

**DOI:** 10.64898/2026.07.03.736426

**Authors:** Robert Pellegrino, Emily J. Mayhew, Jennifer Margolis, Matthew Andres, Alexander B. Wiltschko, Richard C. Gerkin, Joel D. Mainland

## Abstract

Most smells, whether in food, perfume, or the environment, are complex mixtures of many odor molecules. A principled framework for modeling natural mixtures has been considered intractable because mixtures are widely assumed to generate nonlinear perceptual interactions that mirror nonlinear responses at the receptor and neural levels. We quantified perceptual interactions across 432 mixtures composed of 144 component odorants using a trained human panel, and nearly all mixtures were explained by a linear average of their component quality profiles. Within this linear modeling framework, averaging component profiles predicted mixture perception significantly better than the previous state-of-the-art and approached the ceiling set by measurement noise. Even mixtures previously reported to exhibit emergence were equally well predicted. These results suggest that the main challenge in modeling odor perception is characterizing individual components, rather than exhaustively mapping odor–odor interactions. Just as additive color mixing enabled colorimetry, linear mixing in olfaction opens the door to quantitative odorimetry, allowing mixtures to be predicted, reconstructed, and optimized computationally.

## 1 Introduction

The smells we encounter in daily life while eating food or smelling the environment are almost always complex mixtures of many odorants. The olfactory system therefore functions primarily as a detector of mixtures, and predicting how a mixture will smell is a fundamental problem in olfaction. Recent work has made progress on a simpler version of this problem, relating the structure of individual odorants to their perceived qualities (Keller et al. 2017; Lee et al. 2023) and providing a foundation for quantitative models of olfaction. Extending these models to mixtures has been considered more difficult, because mixtures are widely assumed to exhibit qualities not present in the components (emergence) or lose qualities they do carry (suppression) (Dunkel et al. 2014; Thomas-Danguin et al. 2014; Doty 2025; Sisson et al. 2025). These perceptual effects mirror the interactions that mixtures evoke at the receptor and neural level (Pfister et al. 2020; Xu et al. 2020).

The frequency of suppression and emergence in mixtures determines the difficulty of mixture prediction. If suppression and emergence are common, predicting mixture perception requires exhaustively characterizing how its components interact, a combinatorial problem that grows exponentially with the number of molecules. If mixtures are perceived as the average of the component qualities regardless of which components are included (Snitz et al. 2013; Ravia et al. 2020; Dhurandhar et al. 2023), the problem collapses to measuring single-molecule perception for a palette of molecules, which is far more tractable.

Here, we develop a quantitative framework to measure how often odor mixtures exhibit emergent or suppressed qualities. Using descriptive ratings from a trained human panel for 432 mixtures and 144 component odorants, we first asked if the perception of a mixture falls within the range bounded by its components. We then tested how well a mixture’s perception can be predicted by simply averaging the perception of the components, with no interactions between components.

We find that nearly all mixtures smell like a simple average of their components, including those described in the literature as showing emergence or suppression. Averaging component quality profiles predicts mixture perception better than the previous state of the art (Ravia et al. 2020) and approaches the ceiling set by measurement noise. This reframes the main challenge of modeling odor mixture perception as characterizing individual components rather than mapping the vast space of odor–odor interactions. In addition, just as additive mixing turned color into something we could measure and reproduce, linear mixing opens the door to quantitative odorimetry, so we can predict, reconstruct, and optimize mixtures computationally.

## 2 Results

Odor mixtures can be perceived as a blend of their components, or they can exhibit interactions such as suppression or emergence of new perceptual qualities (Figure 1A). To assess how well mixture perception can be predicted from component perception, trained panelists (*n* ≥15, 2 repetitions) evaluated 144 components (115 monomolecular odorants at different concentrations) and 432 mixtures using a 51-descriptor lexicon (Figure 1B, Table S1). This rapid descriptive method, known as rate-all-that-apply (RATA), allowed us to quantify odor characteristics systematically. Our panelists provided stable responses across repetitions (*r*(611) = 0.74, *p <* 0.001; similar to past uses from our lab, Lee et al. 2023; Figure 1C).

**Figure 1:**
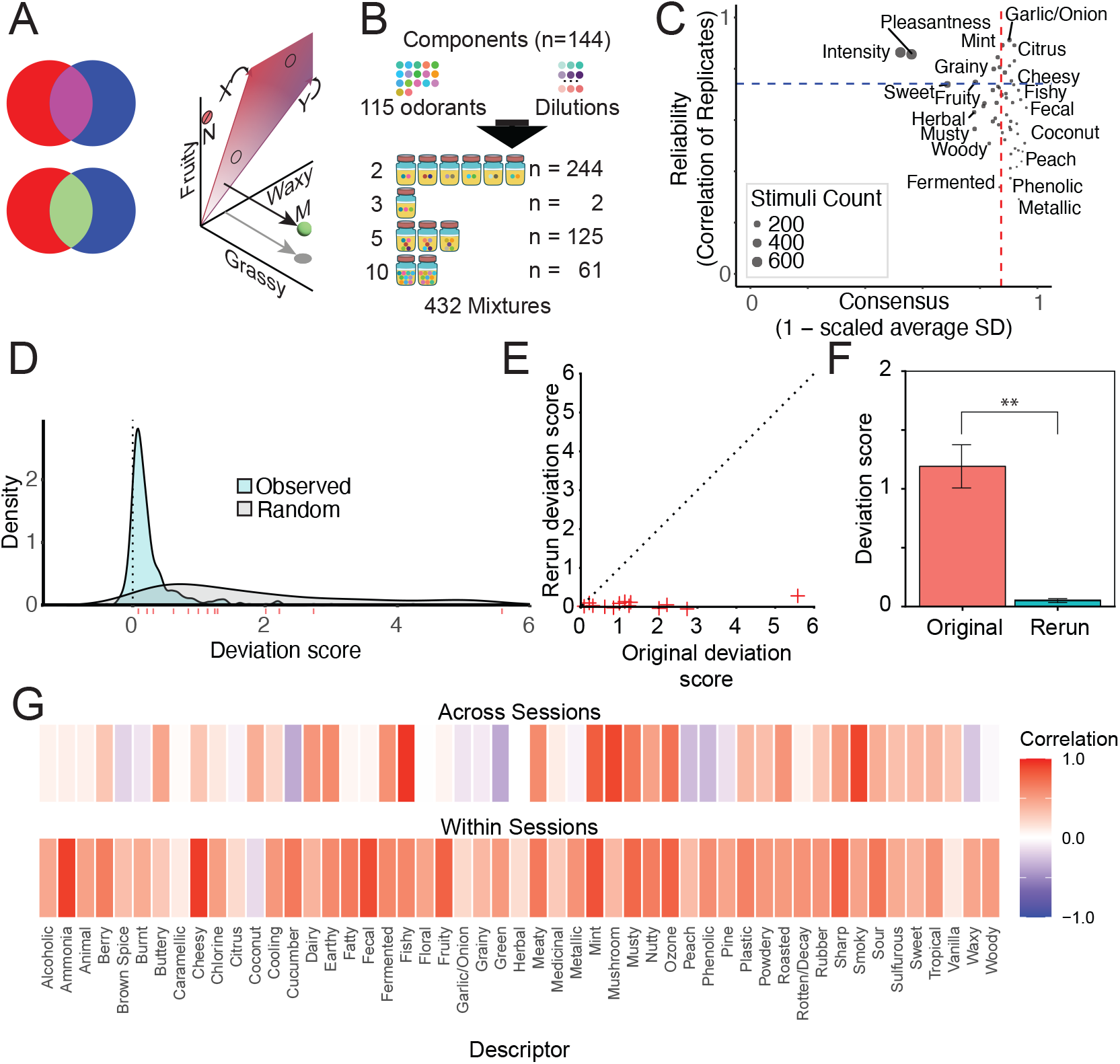
Odor mixture perception is predominantly linear. A) Conceptual illustration of odor mixing in a three-dimensional odor space. Odor mixing may be linear (top color mixing circles, analogous to additive color mixing) or nonlinear (bottom color mixing circles). If mixing is linear, all possible mixtures fall within the shaded region bounded by the component vectors X and Y. In contrast, example mixtures N (suppression) and M (emergence) would be nonlinear. B) 115 component odors at multiple dilutions yielded 144 components, combined into 432 mixtures containing 2–10 components. C) The panelists reliably reported odor quality (*r* = 0.74) and there was a high usage consensus across panelists (0.84). D) Distribution of deviation scores across all mixtures. The deviation score is the cross-replicate dot product of out-of-plane residuals normalized by the in-plane reconstruction (see Methods); zero indicates perfect linearity. The high density near zero indicates that linear mixing dominates. The random distribution shows the same deviation score computed when each mixture is reconstructed from randomly selected components rather than its true components. E) Original versus rerun deviation scores show that controlled within-session replication substantially reduces apparent nonlinearity, consistent with session and panel variation rather than true perceptual interactions. F) Deviation from linearity significantly decreases when mixtures and components are tested within the same session (*p <* 0.003). G) Heatmap of correlations between mixture RATA and averaged component RATA for the 13 re-run samples. Within-session measurements (experiment 2) show stronger consistency than across-session measurements (experiment 1).

### 2.1 Mixture perception is bounded by component perception

To illustrate our test, consider a three-dimensional odor space with only fruity, waxy, and grassy dimensions (Figure 1A). Component X (red) is more fruity than waxy; Component Y (blue) is more waxy than fruity. If mixture perception contains no emergent or suppressed qualities, all mixtures of X and Y should fall in the shaded region. Emergent qualities exceed what any linear combination of components could produce; suppressed qualities fall short. For example, if both components have some waxy character, a purely fruity mixture represents suppression of the waxy character (Fig. 1A, point N). Conversely, if neither component has a grassy character, a grassy mixture indicates emergence of a new quality (Fig. 1A, point M).

Across 432 mixtures spanning the perceptual space, deviation from component-bounded mixing was concentrated near zero (Figure 1D), indicating that emergent and suppressed qualities are rare. We quantified this deviation score by representing each mixture as a 51-dimensional descriptor vector and we used non-negative least squares (NNLS) to find the weighted combination of components that best explains it, with the constraint that no component can negatively contribute to a descriptor. Due to measurement noise, even a perfectly linear system will produce non-zero residuals. To distinguish true nonlinearity from noise, we leveraged the fact that noise-driven residuals should point in random directions across replicates, whereas true nonlinearity should produce residuals that consistently point in the same direction. The deviation score is therefore computed as the dot product of out-of-plane residuals across replicates—positive when residuals point in the same out-of-plane direction across replicates, negative when they point in opposite out-of-plane directions, and near zero when they are uncorrelated—normalized by the in-plane reconstruction, such that a perfectly linear mixture has a score of zero. NNLS treats the 51 descriptors as independent dimensions, so correlated descriptor usage could in principle distort the out-of-plane residual. To rule this out, we repeated the analysis using non-negative matrix factorization (NMF), which re-expresses each mixture in a reduced set of latent factors that capture co-occurring descriptors. Deviation scores remained near zero (Fig. S1), confirming the result is robust across different representations of odor quality.

### 2.2 Session and panel effects account for deviations from component-bounded mixing

Observed deviations from the component-bounded region could stem from true perceptual interactions, experimental noise, or rating variability across panelists and sessions. In experiment 1, inconsistent panel composition correlated with larger deviation scores (Fig. S2). To investigate this further in experiment 2, we retested 13 mixtures spanning the distribution of deviation scores (Table S2), along with their monomolecular components, in a single session with a consistent panel. Deviation scores dropped significantly after this controlled replication (*t*(12) = 3.62, *p <* 0.003; Fig. 1E, F), and within-session correlations were substantially stronger than across-session correlations (Fig. 1G). This suggests that apparent deviations reflect variation in panel composition or temporal drift in ratings across sessions rather than true emergent or suppressive perceptual effects.

### 2.3 Linear averaging approaches the predictive ceiling

In the previous analysis we demonstrated that mixture perception falls within the region bounded by component perception. Here, we examine whether a simple linear model can accurately predict *where* within that region a mixture falls. We extend a previous framework of perceptual odor space for single molecules (Lee et al. 2023) to odorant mixing (Figure 2A). This framework postulates that in perceptual space, the length of the odorant descriptor vector equates to the odorant intensity while the cosine distance to another vector is its quality difference. We extend this framework to postulate that odorants mix within the space of the representable odorant vectors, with the influence on odor quality weighted by their length.

**Figure 2:**
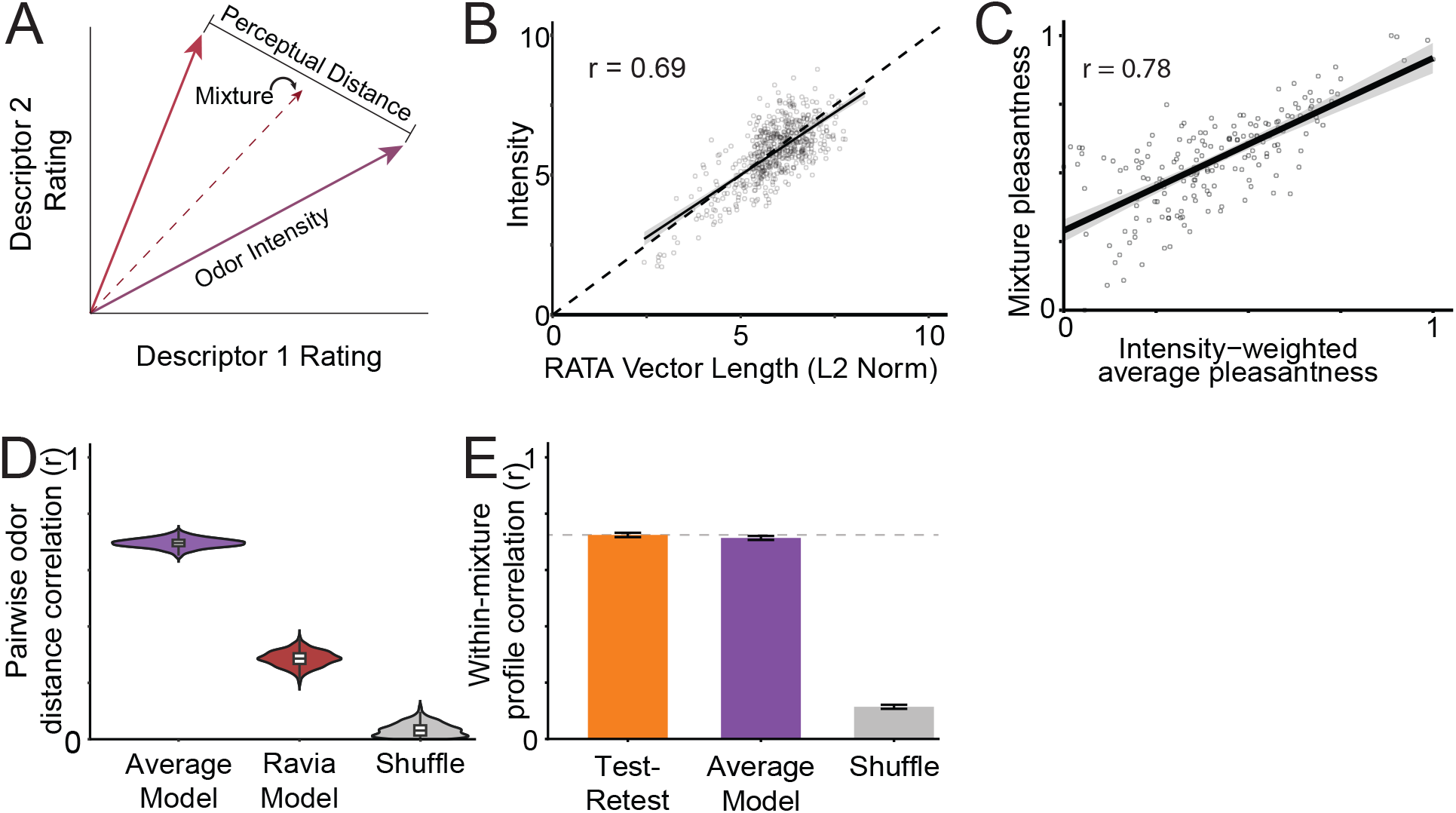
Framework for Odorant Mixing. A) Conceptual extension of perceptual odor space to odorant mixing. Odorant descriptor vectors encode both quality (angle) and intensity (vector length). Mixtures are the average of the component vectors. B) Descriptor vector length correlates with perceived intensity for both single odorants and mixtures. C) Mixture pleasantness shifts systematically with the intensity-weighted average of component pleasantness. D) Correlation of actual and predicted pairwise odor distances (bootstrap distribution over 1000 pairs). The Average Model (purple) significantly outperforms the previous state-of-the-art model (red). E) Per-mixture correlation of the actual and predicted 51-descriptor RATA profile (mean *±* SE) compared to the test-retest correlation between profiles and the correlation between random profiles (grey). The Average Model (purple) performs near the ceiling set by measurement noise (orange reliability ceiling).

We first confirmed that descriptor vector length correlates strongly with perceived intensity (Fig. 2B), both for single odorants (*r*(179) = 0.77, *p <* 0.001) and mixtures (*r*(430) = 0.69, *p <* 0.001), regardless of the number of components (Fig. S3). Mixture pleasantness also follows predictions from this framework, shifting as a function of the intensity-weighted average of component pleasantness (*r*(430) = 0.78, *p <* 0.001; Fig. 2C). We then predicted each mixture’s perception as the unweighted average of its components’ descriptor vectors. This yielded high predictive accuracy (*r*(430) = 0.71, *p <* 0.001), surpassing the previous state-of-the-art model (Ravia et al. 2020; *p <* 0.001, Fig. 2D). To assess whether further improvement is possible on this dataset, we compared model predictions to the ceiling of data reliability, calculated as the correlation between independent replicate measurements of the same mixture. The model was not significantly different from this ceiling (*t*(861) = 0.72, *p* = 0.51; Fig. 2E, S4), suggesting that the remaining error reflects measurement noise rather than model limitations across mixture sizes (Fig. S5).

### 2.4 Even “surprising” mixtures are well predicted by linear averaging

Although the linear averaging model performed well on a large set of mixtures, the literature describes several olfactory mixtures that reportedly exhibit “surprising” effects of suppression or emergence (Sisson et al. 2025). In experiment 3, a human sensory panel with at least 15 subjects performed RATA on eight such mixtures from the literature (Table S3), along with their components (*N* = 26), and reported reference odors (Milo and Grosch 1993; Clery et al. 1999; Le Berre et al. 2008; Sinding et al. 2011; Barkat et al. 2012; Marsili and Laskonis 2014; Sinding et al. 2015). We found that even these “surprising” mixtures did not deviate from linear mixing (Fig. 3A); mixture RATA vectors were highly correlated with model predictions (*r*(406) = 0.84, *p <* 0.001), reaching the ceiling of data reliability (*t*(7) = 1.73, *p* = 0.13, Fig. 3B). For example, one published surprising mixture combines (Z)-4-heptenal (tomato) and (E,Z)-2,6-nonadienal (cucumber) to form a cooked fish odor (Marsili and Laskonis 2014). Although “tomato + cucumber = cooked fish” sounds surprising, this reflects the limitation of representing odor percepts with single words; when profiled via RATA, both component odors have a minor fishy note that stands out in the mixture when other features are reduced through dilution. When the components and mixture are described by the single best descriptor, the predicted and actual vectors are artificially far apart. The similarity improves as more descriptors are used, reaching diminishing returns around five descriptors (Figure 3C).

**Figure 3:**
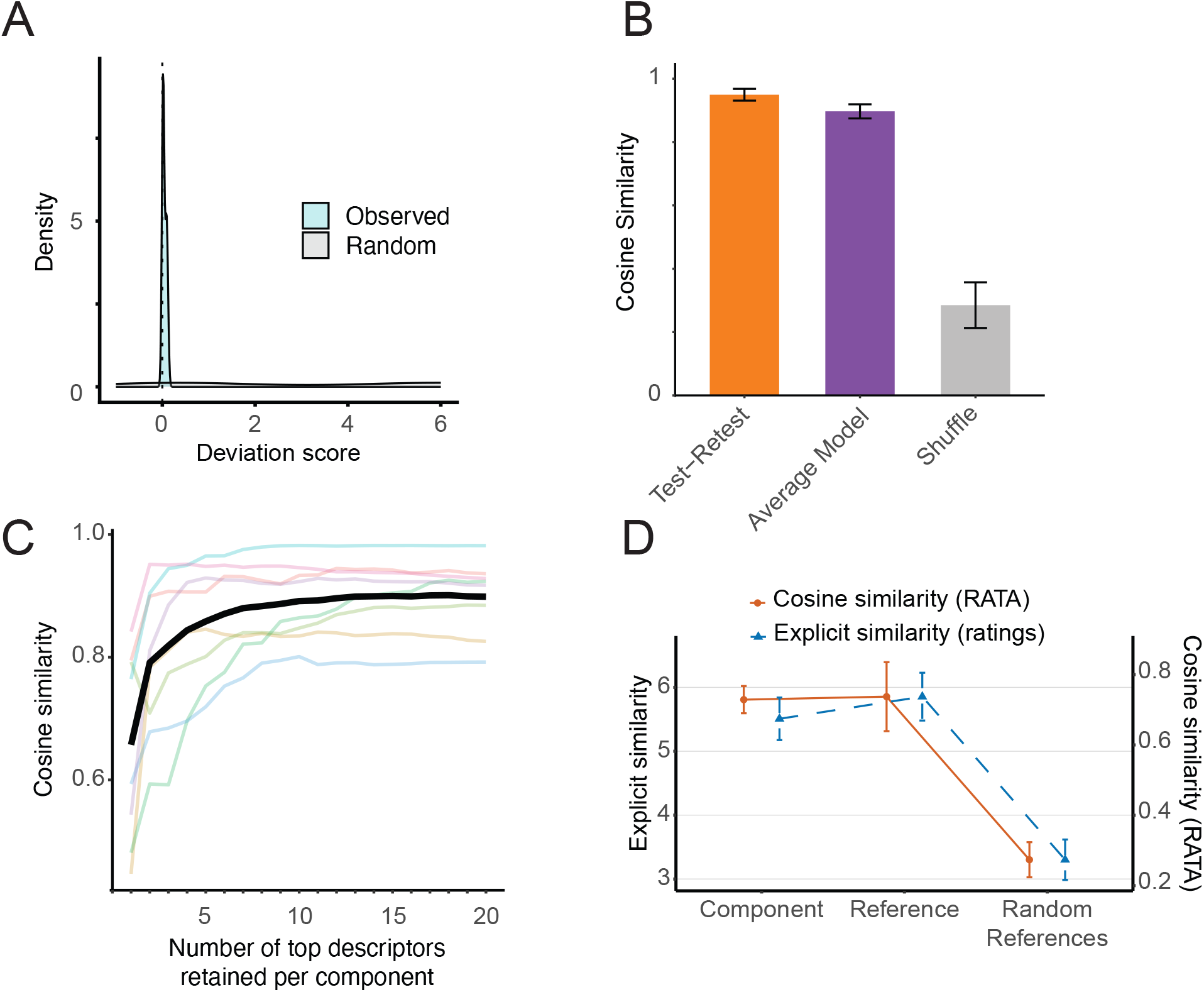
“Surprising” odor mixtures are linear. A) Mixtures previously reported to exhibit suppression or emergence have deviation scores near zero. B) Predictive accuracy of the Average Model for these mixtures (purple) reaches the ceiling of data reliability (orange). C) Descriptor usage in “surprising” mixtures (colored lines; black line, average across all mixtures) is dominated by a small number of descriptors that are consistently among the top descriptors of their components. D) Explicit similarity ratings (blue, left axis) closely track RATA-derived cosine similarity (orange, right axis), with mixtures showing comparable similarity to their components and to reported reference odors, and markedly lower similarity to random references.

One possibility is that our lexicon lacks categories to capture emergent properties. For the example of “strawberry + caramel = pineapple”, the lexicon contains “fruity,” “berry,” and “tropical,” but not “strawberry” or “pineapple.” To test whether our results were an artifact of lexicon limitations, we asked panelists to rate the explicit similarity between components (the strawberry and caramel molecules) and the reference odor (Table S4, allyl hexanoate). Similarity ratings correlated with RATA-derived cosine similarity (*r*(123) = 0.64, *p <* 0.001), validating this approach for discriminating odors. Critically, if mixtures produced emergent qualities distinct from their components, they should be rated as more similar to their reference odors than to their components. Instead, mixtures were rated as equally similar to both (reference: mean (sd) = 5.85 (1.1); components: mean (sd) = 5.51 (1.7), *t*(19) = 0.69, *p* = 0.5, Fig. 3D). Across all three experiments we found no evidence of perceptual interactions between mixture components.

## 3 Discussion

In this study, we found that human perception of odor mixtures can be accurately predicted from component odor perception. Across 432 tested mixtures, the vast majority fell within the perceptual region bounded by their components. Even mixtures previously described as producing emergence or suppression were well described by simple averaging of component profiles. These findings challenge the long-held assumption that olfactory mixture perception is dominated by complex interactions requiring explicit modeling. Instead, they suggest odor perception arises primarily from the properties of individual components rather than emergent mixture effects. This shift in understanding opens the door to simpler models of olfaction, with broad implications for neuroscience, artificial olfactory systems, and fragrance and flavor design.

Although previous studies have reported emergence and suppression in odor mixtures (Laing and Francis 1989; Le Berre et al. 2008; Thomas-Danguin et al. 2014; Sinding et al. 2015; Sisson et al. 2025), many of these reports rely on linguistic labeling or cognitive categorization rather than quantitative perceptual data. When odors are described using single words—such as “pineapple” for a mixture of “strawberry” and “caramel”—apparent emergence may simply reflect the limits of single-descriptor semantic representation. Our multidimensional descriptive approach shows that these mixtures remain predictable from their components across perceptual dimensions, indicating that these phenomena may reflect naming conventions, associative learning, or categorical decision processes, rather than nonlinear sensory interactions at the perceptual level. A second explanation lies in differences in experimental strategy. Many prior studies ask observers to identify individual components within mixtures. As mixture complexity increases, this task becomes more difficult, and components may appear perceptually absent (Laing and Francis 1989). In a linear mixing framework, such effects are expected: adding components reduces the relative salience of any single odor without eliminating its influence on the average. This model is consistent with the concept of olfactory white (Weiss et al. 2012), where multiple random vectors tend to average to the origin.

Our approach differs from previous mixture experiments, which ask whether mixtures are perceived as different from (configural) or the same as (elemental) their components (Kay et al. 2005; Romagny et al. 2018). We instead ask an independent question: whether the descriptive qualities of mixtures are predictable from the descriptive qualities of components. Recognizing the mixture of lime and cinnamon as the configural percept of cola, rather than as two distinct elements, likely depends on prior exposure to cola. Regardless of whether it is perceived configurally or elementally, we hold that the quality profile can be predicted by averaging the component qualities.

Mechanistically, nonlinear interactions exist at the receptor and neural level (Singh et al. 2019; de March et al. 2020; Pfister et al. 2020; Xu et al. 2020; Zak et al. 2020), but it remains unclear how many of these interactions occur within the typical intensity ranges of natural odors (Wachowiak et al. 2025). Regardless, mechanistic nonlinearity does not imply perceptual nonlinearity. The visual system illustrates this clearly: despite numerous nonlinearities at the receptor and neural level, perceptual color mixing is predominantly linear (Stockman and Brainard 2010). How nonlinear neural representations give rise to effectively linear perceptual spaces remains an important open question.

Our findings should be interpreted in light of several limitations. First, if an emergent quality cannot be properly captured by the available lexicon, the mixture may appear artificially well predicted. Second, measurement noise—from panel variability, session effects, and perceptual drift— introduces uncertainty that can obscure deviations from our model. Third, our stimulus set was not fully representative of natural olfactory conditions. All components were diluted to moderate intensities, and mixtures contained no more than ten components, whereas natural odors often contain hundreds of components with individual intensities spanning a much broader range including below detectable levels. Fourth, odor perception can change with concentration (Gross-Isseroff and Lancet 1988; Laing et al. 2003), which contrasts with our averaging framework where an odorant mixed with itself should produce the same perceptual profile regardless of concentration. We predict that we did not see this effect because the concentration of a component alone and in the mixture differed by less than an order of magnitude in all experiments. While accounting for quality changes across concentration is important for understanding odor coding mechanisms, it may be less relevant in practice: most applications involve formulating mixtures from characterized components at fixed concentrations, rather than predicting perception across large concentration changes.

The predominance of linearity in mixture perception has several important implications. First, it suggests that olfactory metamers, distinct mixtures that smell identical, are not rare but an inherent consequence of averaging, since many different component combinations can produce the same perceptual vector. Second, this framework supports the pursuit of *primary odors*: a minimal set of basis stimuli that, through linear combination, could reconstruct the full perceptual space of smell. Finally, just as the discovery of additive color mixing enabled colorimetry, a quantitative system for describing and predicting color, olfaction may be amenable to a parallel framework of quantitative odorimetry, where mixtures can be numerically represented. This shift reframes olfaction from a system seemingly dominated by unpredictable interactions to one that can be mapped, predicted, and synthesized using linear principles.

## 4 Methods

### 4.1 Participants

A total of 30 prospective panelists (ages 18–55, normal olfactory function) were recruited from the Philadelphia area. All participants provided informed consent in accordance with the protocol approved by the University of Pennsylvania Institutional Review Board (protocol #843955). Prior to study onset, all participants were screened for their ability to smell and describe odors reliably with scales (see Training and Screening).

### 4.2 Lexicon

We selected 51 of the 138 common labels from a merged Goodscents–Leffingwell dataset (Lee et al. 2023) to form our lexicon. Selection prioritized broad category terms (e.g., fruity, floral) to span the range of possible odor percepts, while also including a limited set of specific terms within categories to measure model precision (e.g., citrus, berry). We additionally included negative odor terms absent from the 55-term lexicon used in prior data collection from our lab (Lee et al. 2023) and often absent from perfumery datasets. Each lexicon term was grounded to a physical reference to promote agreement among panelists. The final list of descriptors and associated references is provided in supplementary data (Table S1).

### 4.3 Training and Screening

Sessions 1–5 constituted a training and screening protocol. In Session 1, panelists were introduced to the rate-all-that-apply (RATA) method (Dravnieks 1985; Meyners et al. 2016), in which descriptive terms are selected from a lexicon and rated on a 0–5 scale for applicability, and completed a pre-test by evaluating 20 common odorants. Sessions 2 and 3 focused on lexicon training: researchers defined each odor term, provided visual aids, and guided panelists in smelling corresponding odor references; quizzes with blinded odorants and mixtures reinforced learning. In Sessions 4 and 5, panelists re-evaluated the 20 common odorants. Performance was assessed by test–retest reliability and appropriateness of label use. Panelists who met inclusion criteria (individual test-retest correlation *R >* 0.35; reasonable label assignment for common odorants) advanced to the sensory panel.

### 4.4 Experiment 1

At least 15 trained panelists evaluated each of 144 components (monomolecular odorants diluted to a certain concentration) and 432 odor mixtures using the 51-descriptor RATA lexicon across 16 sessions. Odorants were selected to span molecular space, perceptual space, and a range of intensities (Figure S6); mixing was pseudorandom.

### 4.5 Experiment 2

To assess whether variability in panel composition or rating drift across sessions influenced Experiment 1 results, we selected 13 mixtures representing cases of low, moderate, and high deviation scores (Table S2). In a single session, a consistent panel profiled these 13 mixtures and all of their components. The RATA methodology was identical to prior sessions.

### 4.6 Experiment 3

Several odor mixtures have been reported to evoke percepts unlike their components in the literature (Milo and Grosch 1993; Clery et al. 1999; Le Berre et al. 2008; Sinding et al. 2011, 2015; Barkat et al. 2012; Marsili and Laskonis 2014). To further test the assumption of linearity, we measured RATA responses for eight of these mixtures, their 26 component odorants at concentrations reported in the literature, and representative references (Table S3, S4) in the same session and using a consistent panel.

### 4.7 Perceptual Similarity

Distances between two odors measured using RATA will vary based on the lexicon. To test if the perceptual distances were dependent on our lexicon, we repeated a subset of measurements using explicit similarity (Snitz et al. 2013; Ravia et al. 2020). Panelists were first asked to rate stimuli that were identical, similar (“lime” vs. “lemon”), or perceptually distant (“garlic/sulfur” vs. “pineapple”) to help calibrate their scaling. Next, panelists reported pairwise similarity for surprising odors, their components, and the relevant reference in two duplicate sessions.

### 4.8 Data Analysis

Of the original 54 rated terms, “intensity”, “pleasantness” and “no odor” were removed prior to analysis, leaving 51 terms describing odor character. All analyses used stimulus–annotator combinations with exactly two replicates (*<* 0.1% excluded). For each mixture–component comparison, only data from annotators who had described both the mixture and its components were retained, and only when at least 5 such annotators were available, to reduce spurious effects from distinct annotation pools. The resulting dataset thus contains, for each stimulus, 51-dimensional descriptor vectors from 5+ annotators in duplicate. The mean 51-dimensional vector across annotators for a single replicate constitutes the “response” to that stimulus.

To characterize panel performance, we computed two metrics for each of the 51 descriptors. Reliability was assessed as the Pearson correlation between replicate-averaged stimulus vectors (rep 1 vs. rep 2), computed across all stimuli for each descriptor, then summarized as the mean correlation across stimuli and descriptors. Consensus was defined as 1*−* (mean within-descriptor SD*/*SD_max_), where SD was computed across annotators for each stimulus before averaging across stimuli; lower inter-annotator variability thus yields higher consensus.

To test whether each mixture’s response could be predicted by a weighted sum of its components’ RATA values, we used non-negative least squares (NNLS) regression. NNLS estimates how much each component contributed to the overall mixture response (i.e. addition in plane), and how much of the response could not be explained by a (non-negatively) weighted sum (i.e. residual out of plane); NNLS finds the set of weights that minimize this out of plane residual, given the constraint that weights cannot be negative (i.e. a mixture component that is “fruity” cannot negatively contribute to the fruitiness of the mixture). The out-of-plane residual vector is simply the mixture response vector minus the in-plane sum obtained from the NNLS solution, and its direction and magnitude capture deviation from linearity.

Due to measurement noise, this out-of-plane residual will be non-zero even for a perfectly linear system. However, if deviations arise purely from noise, their direction should be random across replicates. We therefore computed the out-of-plane residual for each replicate and took their dot product: positive values indicate correlated (consistent) deviations, negative values indicate anti-correlated deviations, and values near zero indicate the noise-driven uncorrelated deviations expected under linearity. We further computed a deviation score—the relative magnitude of out-of-plane (dot product of residual of first replicate with that of second replicate) to in-plane contributions (dot product of in-plane reconstruction of first replicate with that of second replicate), as a summary measure of the fraction of odor character deviating from linearity. Under the null hypothesis of linearity, this ratio should be zero, meaning that mixture responses are fully accounted for as non-negative weighted sums of the component responses.

For each two-component mixture, intensity and pleasantness ratings were averaged across annotators and replicates. Mixture pleasantness was then predicted as the intensity-weighted mean of component pleasantness: (*P*_1_ *· I*_1_ + *P*_2_ *· I*_2_)*/*(*I*_1_ + *I*_2_), where *P* and *I* denote pleasantness and intensity ratings for each component. Predictive accuracy was evaluated by Pearson correlation between predicted and observed mixture pleasantness across all two-component mixtures.

For surprising mixtures, to assess how much of a mixture’s perceptual character is carried by its components’ dominant descriptors, we varied the number of top-rated descriptors *k* (1–51) retained per component. For each *k*, all but the *k* highest-rated descriptor values in each component vector were set to zero, the zeroed vectors were averaged across components to form a sparse linear prediction, and cosine similarity between this prediction and the observed mixture vector was computed. This was repeated for each mixture and its trajectories are shown in Figure 3C.

Finally, we compared the predictive performance to the current state-of-the-art (SOTA) angle distance model (Ravia et al. 2020) for pairwise mixture comparisons in the dataset from experiment 1 (*n* = 26,335). Physicochemical distances between mixtures were computed following Ravia et al. (2020). Each mixture was represented as the summed Dragon descriptor vector of its constituent molecules (21 features selected by Ravia et al.), and pairwise angular distances between mixture representations were calculated. Perceptual distances between the same mixture pairs were computed as the angle between their mean 51-dimensional descriptor vectors. Pearson correlation between physicochemical and perceptual distances was computed over 1,000 randomly sampled mixture pairs; bootstrap confidence intervals were obtained by resampling with replacement (1,000 iterations). This Ravia-model correlation was compared against the linear (intensity-weighted average) model correlation using the same 1,000 pairs.

## Acknowledgements

This research was supported by Google Research and National Institutes of Health grants F32DC020380 (to RP), R01DC017757, and U19NS112953. We thank Brittney B. Nguyen, Tanushri Bhatnagar, Carissa Evans, and M. Dougherty for their help designing stimuli.

## Author contributions

RP, EJM, ABW, RCG, and JDM conceived of the experiment(s). RP and RCG wrote the analysis code and analyzed the data. JDM, JM, MA, and EJM collected the data. RP and JDM wrote the manuscript, and all authors edited the manuscript.

