## Supplementary Figures for "Odors Smell Like Their Components: A Linear Framework for Predicting Olfactory Mixture Perception"

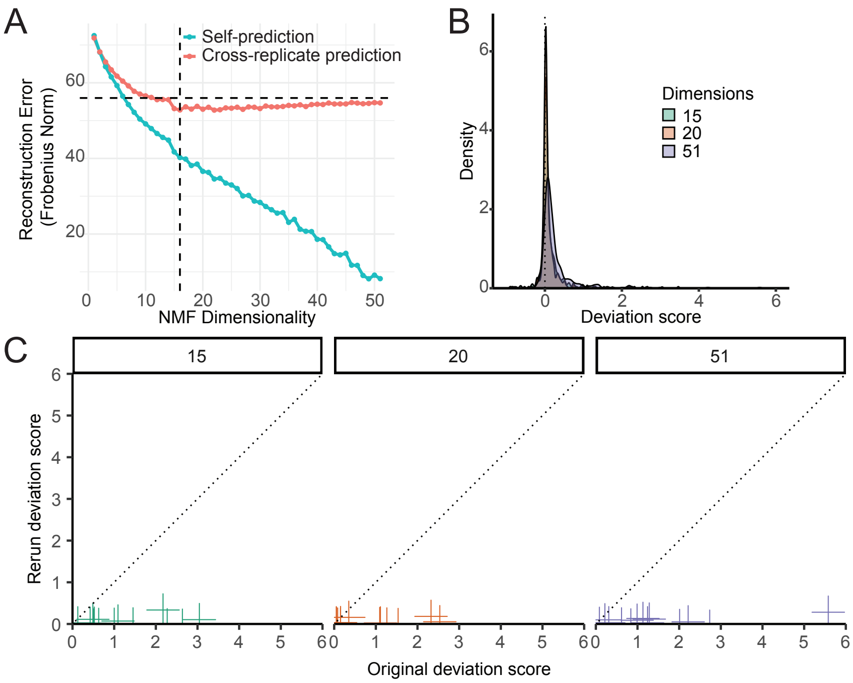


**Figure S1: Determination of NMF dimensionality and its effect on deviation scores.** A) Reconstruction error (Frobenius norm) as a function of the number of NMF dimensions. When a replicate is reconstructed from a compressed version of *itself*, error decreases monotonically with added dimensions (teal). When a replicate is reconstructed from a compressed version of an *independent* replicate, error plateaus at 15 dimensions (red; dashed lines), indicating that dimensions beyond 15 capture replicate-specific noise rather than shared structure. B) Distribution of deviation scores computed at 15, 20, and 51 (full descriptor set) dimensions. C) Original versus rerun deviation scores at each dimensionality. Reducing dimensionality from 51 to 15 does not change the overall conclusion that most mixtures cluster near the origin, indicating predominantly linear mixing.

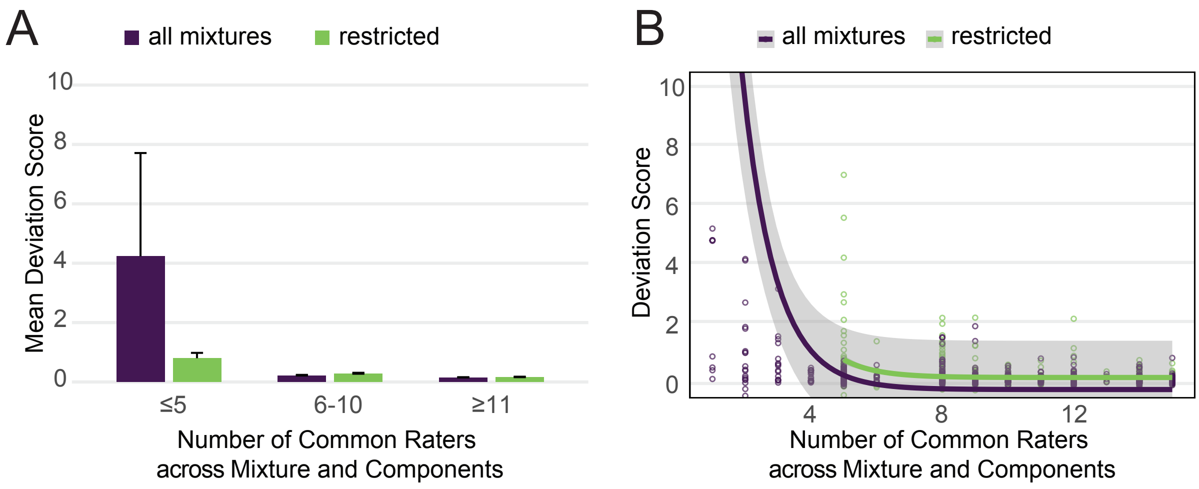


**Figure S2: Rater overlap and number of raters influence apparent deviation from linearity.** A) Mean deviation from linearity as a function of the number of raters common to a mixture and its components, binned as $\leq 5$, 6–10 and $\geq 11$. "All mixtures" includes every mixture; "restricted" retains only mixtures and components rated by the same panelists and excludes mixtures with fewer than five common raters. Deviation scores are substantially higher when fewer raters scored both mixture and components, but this effect diminishes as the number of common raters increases. B) Deviation score versus the ungrouped number of common raters, with fitted trend lines (shaded regions, 95% confidence intervals) for the "all mixtures" and "restricted" samples, showing the same convergence at higher rater counts.

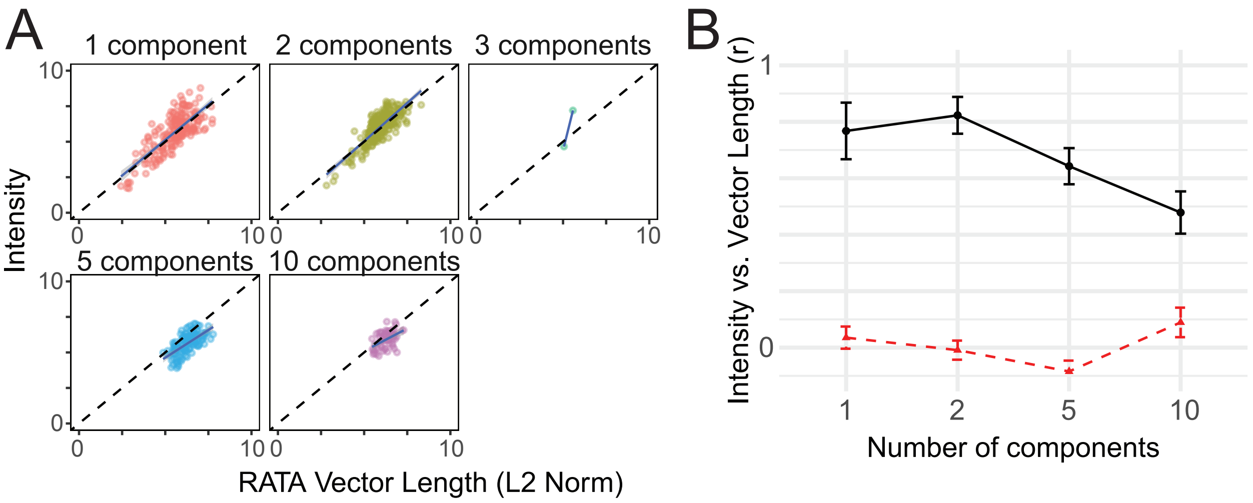


**Figure S3: RATA vector length correlates with intensity across mixture sizes.** A) Unit-slope plots relating descriptor vector length (L2 norm) to perceived intensity, shown separately for stimuli with 1, 2, 3, 5, and 10 components. B) Correlation ($r$) between vector length and intensity for stimuli with 1, 2, 5, and 10 components (black; mean $\pm$ SE). The red dashed line shows the correlation after shuffling intensity values within each mixture size, confirming that the observed relationship is not attributable to chance.

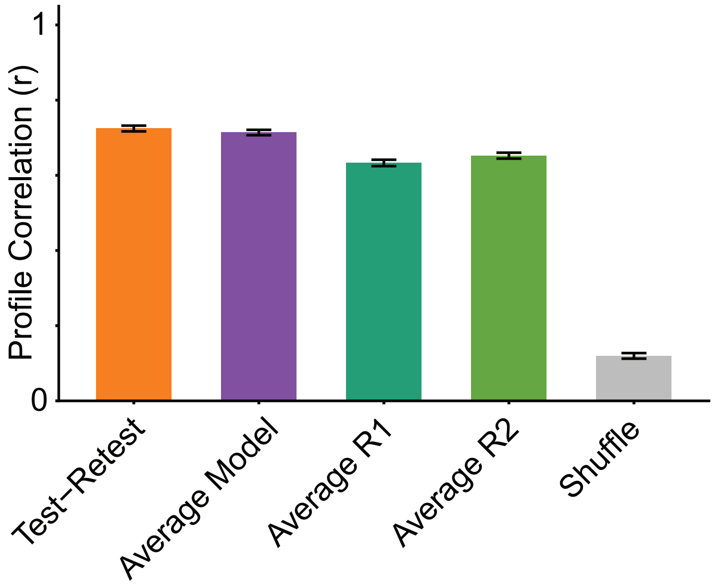


**Figure S4: Model performance approaches the test-retest reliability ceiling.** Correlation (Pearson $r$) between predicted and actual 51-descriptor RATA profiles, pooled across all mixtures and descriptors, under five comparison schemes. Test-Retest reflects the correlation between mixture RATA replicate 1 versus replicate 2 ($r = 0.73$). Average Model uses component RATA averaged across both replicates to predict averaged mixture RATA ($r = 0.71$), performing comparably to Test-Retest ($p = 0.51$). Average R1 uses replicate 1 component RATA to predict replicate 2 mixture RATA ($r = 0.62$), and Average R2 uses replicate 2 component RATA to predict replicate 1 mixture RATA ($r = 0.64$); both fall below Test-retest once the averaging advantage is removed ($p < 0.001$ for both). Shuffle reflects the correlation between randomly paired profiles ($r = 0.14$), significantly lower than both the Average Model and Test-Retest ($p < 0.001$ for both).

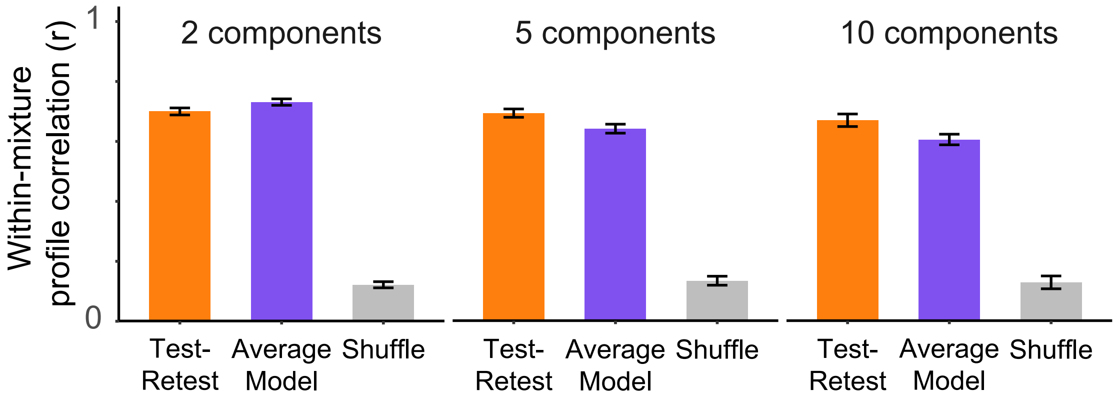


**Figure S5: Model performance across number of mixture components.** Per-mixture correlation (Pearson $r$) between the actual and predicted 51-descriptor RATA profiles, shown separately for 2-, 5-, and 10-component mixtures, alongside the test-retest correlation between replicate profiles and the correlation between randomly paired (shuffle) profiles. The Average Model performs near the test-retest ceiling and far above chance at every mixture size.

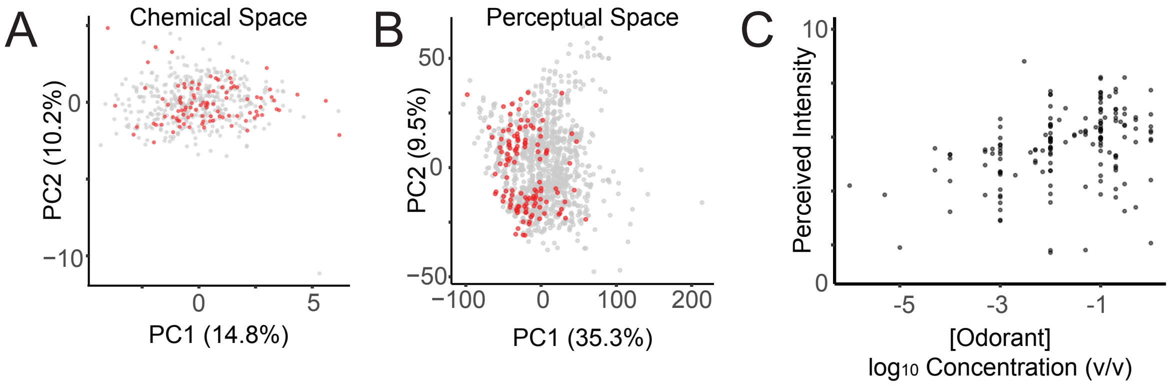


**Figure S6: Selected odorants span molecular space, perceptual space, and a range of intensities.** A) Chemical space, visualized by principal component analysis (PCA) of Dragon molecular descriptors across all candidate odorants (gray); odorants selected for this study (red) span a broad region. B) Perceptual space, visualized by PCA of panel-averaged RATA profiles from the DREAM Olfaction Challenge 2025 dataset across all odorants (gray); selected odorants (red) similarly span a broad region. C) Perceived intensity as a function of concentration (v/v, log scale), showing that selected odorants were tested across a wide range of concentrations and resulting intensities.
